# Bone Morphogenetic Protein (BMP) signaling regulates germline stem cell self-renewal in the newly formed *Drosophila* testis stem cell niche

**DOI:** 10.1101/2022.03.26.485802

**Authors:** Merci N Best, Ashley Fidler, Marieke K Jones, Matthew Wawersik

**Author notes:** Corresponding author (MW). Department of Biology, University of Virginia, Charlottesville, Virginia, United States of America (Merci N Best). Department of Chemistry, Princeton University, Princeton, New Jersey, United States of America (Ashley Fidler). These authors contributed equally to this work (Merci N Best, Ashley Fidler).

## Abstract

Bone morphogenetic proteins (BMPs) are a group of multifunctional cytokines and metabologens highly conserved in the transforming growth factor-β (TGF-β) superfamily. BMPs have an established role in controlling cell fate and tissue morphogenesis in a diverse range of organisms and are known to maintain stem cells in adult *Drosophila* testes. Additional studies in embryonic and larval testes suggest BMP regulates germ cell behavior. However, the roles of BMP signaling in controlling testis stem cell formation have yet to be directly examined. Here, we explore the pattern of BMP activation during embryonic testis niche morphogenesis as well as niche maturation in larval testes. We also assess the impact of altered BMP signaling on these stages of development. Our data suggest that BMP signaling is critical for promoting germ cell identity in primordial germ cells during embryonic niche formation. During niche maturation, we also find that that BMP signaling is both necessary and sufficient for maintenance of a self-renewing germline stem cell (GSC) population, and that newly formed cyst stem cells (CySCs) are a source of BMP activating ligand. As development progresses, however, BMP activation is no longer sufficient to alter GSC self-renewal. Collectively, our work promotes a more thorough understanding of BMP as a key mechanism controlling stem cell development in *Drosophila* testes that has implications for the development of other organ systems.

## Introduction

The Transcription Growth Factor-β/Bone Morphogenetic Protein (TGF-β/BMP) signaling pathway is linked to developmental disorders, cancer, and senescence [1–11]. BMPs have established roles controlling cellular proliferation, differentiation, and apoptosis in mammalian tissues, including female and male reproductive organs [12–17]. The BMP pathway also controls cell fate and tissue morphogenesis in organisms ranging from humans to the fruit fly, *Drosophila melanogaster* [18–24].

The *Drosophila* BMP pathway is activated by one of three secreted ligands: Decapentaplegic (Dpp), Glass-bottom-boat (Gbb) or Screw (Scw) (Fig 1A; see [29, 31] for comprehensive reviews). In the canonical pathway, these ligands bind cell surface receptors that form via heterodimerization of the type I and type II transmembrane serine/threonine kinases, which include Thickveins (Tkv), Saxophone (Sax) or Punt [32, 33]. Ligand binding induces phosphorylation of the type II receptor, Tkv, by either Sax or Punt which leads to phosphorylation of the SMAD protein, Mothers-against Decapentaplegic (Mad), by Tkv. Subsequently, phosphorylated Mad (pMad) complexes with the co-SMAD, Media (Med), and enters the nucleus where it functions as a transcription factor to modulate gene expression.

**Fig 1.**
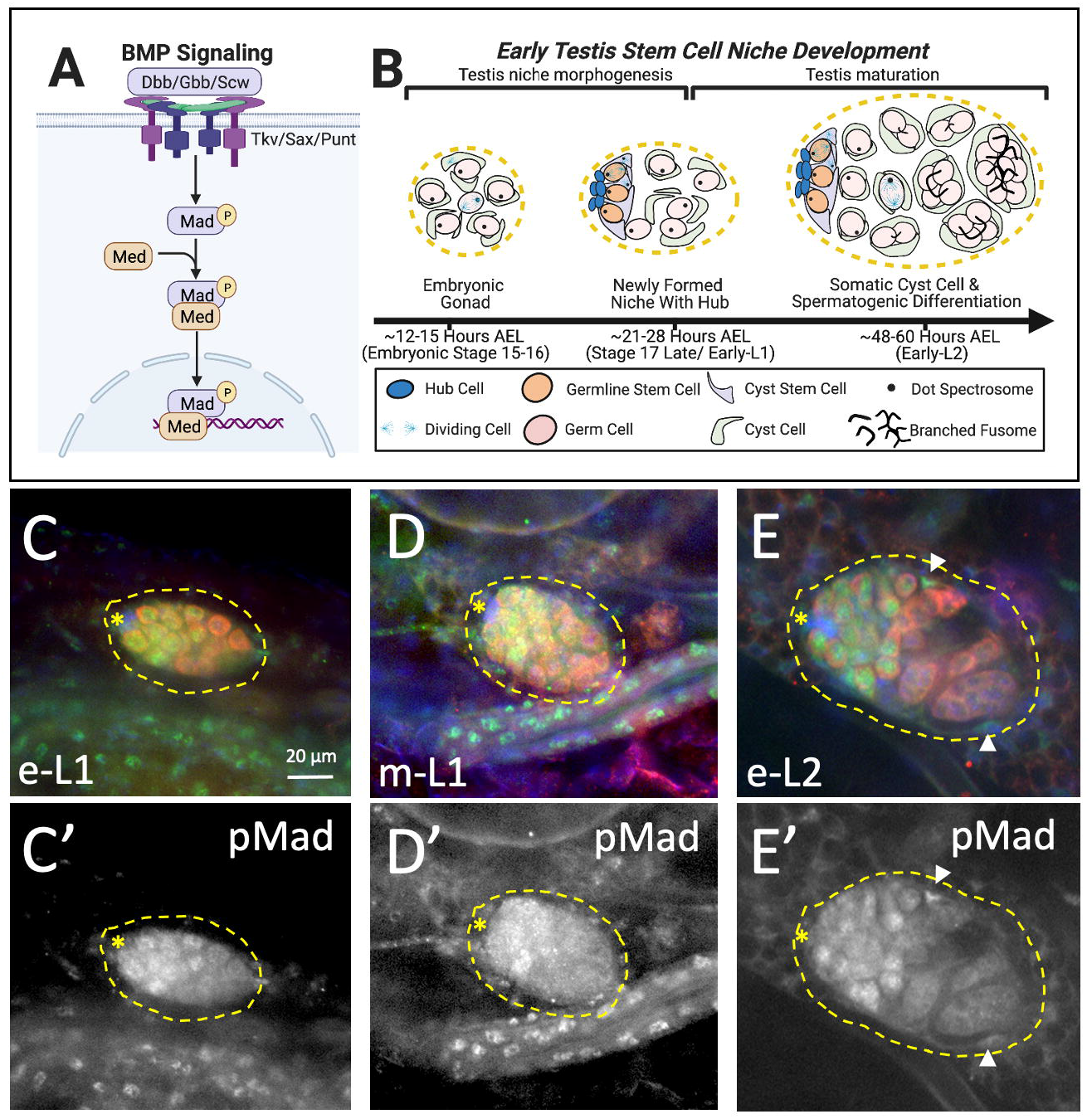
BMP signaling is activated during maturation of the testis GSC niche. (A) Schematic of the canonical Bone Morphogenetic Protein (BMP) signaling pathway resulting in nuclear localization of phospho-activated Mothers Against Decapentaplegic (pMad) in complex with the co-transcriptional activator Media (Med). (B) Schematic of early testis stem cell niche development showing morphogenesis of a functional niche with GSCs and CySCs docked at a tightly coalesced hub by the embryo to larval transition (∼22 hrs AEL), followed by maturation of the testes resulting in onset of spermatogonial and cyst cell differentiation in mid-L1 testes (32-40 hrs AEL; not shown) and an abundance of differentiating spermatogonia with elongated and branched fusomes in the testes posterior by early-L2 (∼48 AEL). (C-E) Immunostaining of wild type larval testes for BMP activity as assessed by pMad (green or alone in grayscale; C’-E’) co-stained with Vasa (red) to detect germ cells and Fas3 (blue) to detect the hub. Testis outlined with asterisk indicating location of the hub. Scale bar is 20 µm. Testes from (C, C’) early-L1 (n=14), (D, D’) mid-L1 (n=19), and (E, E’) early-L2 (n=11) show nuclear pMad staining in germ cells at the testis anterior where GSCs and gonialblasts reside. pMad is also detected in somatic cells in the posterior half of early-L2 testes (E, E’; arrow heads) where differentiating cyst cells reside. Panel A created through BioRender.com.

*Drosophila* testes have proved an important system for understanding how BMP regulates cell behavior [34, 35]. Adult male flies contain a pair of testes; each of which forms a coiled tube that houses the germline stem cell (GSC) niche at its apex [35]. This niche functions to ensure continuous gamete production in sexually reproducing organisms [34, 36]. Three distinct cell types are associated with the testis niche: sperm producing GSCs, somatically derived cyst stem cells (CySCs), and somatic hub cells. Hub cells reside at the tip of each testis where they form a tightly adherent cluster of 10-15 quiescent somatic cells that regulate the division and differentiation of GSCs and CySCs [34, 37]. Specifically, 5-9 undifferentiated GSCs are arrayed around the hub where they can divide asymmetrically to produce two daughter cells: one GSC that remains docked at the hub, and one gonialblast daughter that is displaced from the hub [38]. Continued gonialblast divisions produce a 16-cell spermatogonia that further differentiates and initiates meiosis to produce 64 individualized sperm [34, 36].

Like the hub cells, CySCs are critical for controlling GSC maintenance and differentiation (see [34,39,40] for comprehensive reviews). An array of 15 - 20 somatic CySCs form membrane projections that surround the self-renewing GSCs and enable CySC-hub adhesion [34, 41]. Asymmetric division of CySCs localized to the hub produce one self-renewing CySC that remains associated with the hub and GSCs, and a second daughter cyst cell that functions to nurture spermatogenic differentiation [42]. Pairs of cyst cell daughters ensheath newly-produced gonialblast cells as they are displaced from the hub [43]. The cyst cell pairs, which cease to divide, then grow in size to encase the differentiating germ cells as they proceed through spermatogenesis [41, 44].

BMP signaling in the adult testes niche is essential for GSC self-renewal. Hub cells and CySCs secrete Dpp and Gbb to activate BMP signaling in neighboring GSCs [35, 45]. In turn, BMP activation represses GSC differentiation via repression of BAM expression, while gonialblast cells displaced from the niche are permitted to differentiate [15,45–48]. Although much is known about BMP regulation of GSC maintenance and differentiation in the adult, less is known about BMP function during testis development.

In developing *Drosophila*, testes form when primordial germ cells (PGCs) from the posterior of the embryo associate with surrounding somatic gonadal precursor (SGP) cells [49–51] (for comprehensive reviews see – [19,20,52]). PGCs migrate from the embryo posterior toward a cluster of SGPs that form bilaterally within the embryonic mesoderm, after which these two cell types coalesce into a compacted gonad by the middle of embryogenesis at ∼10.5 hrs after egg laying (AEL) [19, 20]. Just prior to gonad coalescence, hub cells are specified from a subset of anterior SGPs via signaling from the overlying midgut [53, 54]. After coalescence, niche morphogenesis results in formation of a functional GSC niche with GSCs and CySCs arrayed around a hub by the end of embryogenesis at ∼22 hrs AEL (Fig 1B) [53–58]. During early larval development, testes continue to mature, with GSCs and CySCs initiating asymmetric division to produce functional cyst cells that nurture spermatogonial maturation in the testis posterior by mid-L1 (∼32 hrs AEL) [56–58].

The BMP signaling pathway regulates germ cell behavior in specific stages of testes development. BMP signaling is activated in germ cells at the posterior pole of the embryo prior to migration [19,59,60], and BMP promotes PGC migration toward SGPs [27]. Work by Deshpande et al (2014) also suggests that BMP activation is required for maintenance of PGC identity during early stages of embryonic gonad formation [27]. Finally, Chang et al (2013) have shown BMP signaling is strongly activated in GSCs of L1 and L2 larvae during early stages of testis niche morphogenesis, but this activity is downregulated in L3 larval (72-120 hrs AEL) and adult testes by the E3-ubiquitin ligase, Smurf. In this study, Smurf was found to control larval testes growth, GSC number, and spermatogonia division [61]. While Smurf has been shown to control stem cell behavior during larval testis maturation, the role of BMP regulation has not been directly examined at this time. Additionally, BMP function during testis niche morphogenesis in late-stage embryos has not been studied. Work described here strives to bridge this knowledge gap through a direct analysis of BMP activation and function during testis niche morphogenesis and maturation.

## Results

### BMP signaling is activated during testis niche maturation

To better understand the role of BMP during testis development, we first sought to determine the pattern of BMP activation during testis niche maturation (Fig 1). We, therefore, utilized a previously characterized phospho-Mad (pMad) antibody [62–64] to examine BMP activity in wild type larval testes after a functional testis niche has formed. We find that pMad is initially detected in the nuclei of germ cells in early-L1 and early/mid-L1 testes (24-28 and 28-32 hrs AEL), with expression biased toward the testes anterior where GSCs and CySCs have just formed (Figs 1C and S1). As testes mature to produce differentiating spermatogonia and cyst cells in the testis posterior, the pMad staining pattern becomes further restricted to the nuclei of GSCs and early gonialblasts in mid-L1 testes (32-40 hrs AEL), with only diffuse cytosolic staining observed in the testis posterior (Figs 1D and S1). This GSC-biased staining pattern persists in late-L1/early L2 larval testes (40-60 hrs AEL, where nuclear pMad is also detectable in differentiating somatic cyst cells (Figs 1E, S1, and S2). This pattern is similar to data previously obtained in 1^st^ instar larval testes [61], and is consistent with a role for BMP signaling in the regulation of GSC self-renewal in the newly formed niche comparable to adult testes.

### BMP signaling controls germline stem cell maintenance during early niche maturation

To directly assess whether BMP activation in the newly formed testes niche is responsible for controlling GSC self-renewal, we assessed whether BMP signaling is necessary and sufficient for maintenance of undifferentiated GSCs in early larval testes. A key marker of GSC differentiation status is the fusome; a germline-specific organelle that functions in microtubule organization, signaling, and cellular transport (schematic in Fig 1B; [65, 66]). In GSCs and gonialblast cells, fusomes appear as a small, spherical “dot” in the cytoplasm that is also called a spectrosome. During germ cell differentiation, the fusome elongates to form a “bar” structure that spans the interconnected cytoplasm of 2-cell spermatogonia, and then grows into a “branched” morphology that extends between 4-, 8- and 16- cell spermatogonia (Fig 1B).

To assess whether BMP signaling is necessary for GSC maintenance in newly formed testes, we therefore examined the impact of BMP inhibition on fusome morphology (Figs 2A-C and S3). Specifically, the Gal4-UAS system [67] was used to ectopically express inhibitors of BMP signaling in developing germ cells. As spermatogenic differentiation marked by fusome branching is first observed in the latter half of the larval 1st instar [57], testes in late-L1/early-L2 larvae (40-60 hrs AEL) were examined. Germline over-expression of Daughters against decapentaplegic (Dad), a feedback inhibitor of cytosolic BMP signaling [68], results in testes showing elongated and branched fusomes in germ cells adjacent to the hub (Fig 2B). This was observed in 100% of testes after ectopic Dad expression, while 0% of y,w1118 control testes showed this phenotype (S3 Fig). Consistent with a defect in maintenance of self-renewing GSCs, germline Dad over-expression also resulted in an ∼30% reduction in testes size on average (S.D.±2.7, P-value = 6.16 x10^-5^) compared to controls (Fig 2A-B). Similar observations were made after germline over-expression of a dominant negative allele for the type II BMP receptor, Tkv (Tkv^DN^; [69]). Once again, 100% of Tkv^DN^ over-expressing testes showed elongated or branched fusomes in germ cells near the hub (Figs 2C and S3). Additionally, ectopic Tkv^DN^ expression in germ cells results in a reduction in testes size by ∼17% on average (SD±2.7, P-value = 2.1×10^-4^) compared to controls (Fig 2A, C). Together, these data indicate that BMP signaling in the germline is necessary for GSC maintenance during early stages of larval niche maturation.

**Fig 2.**
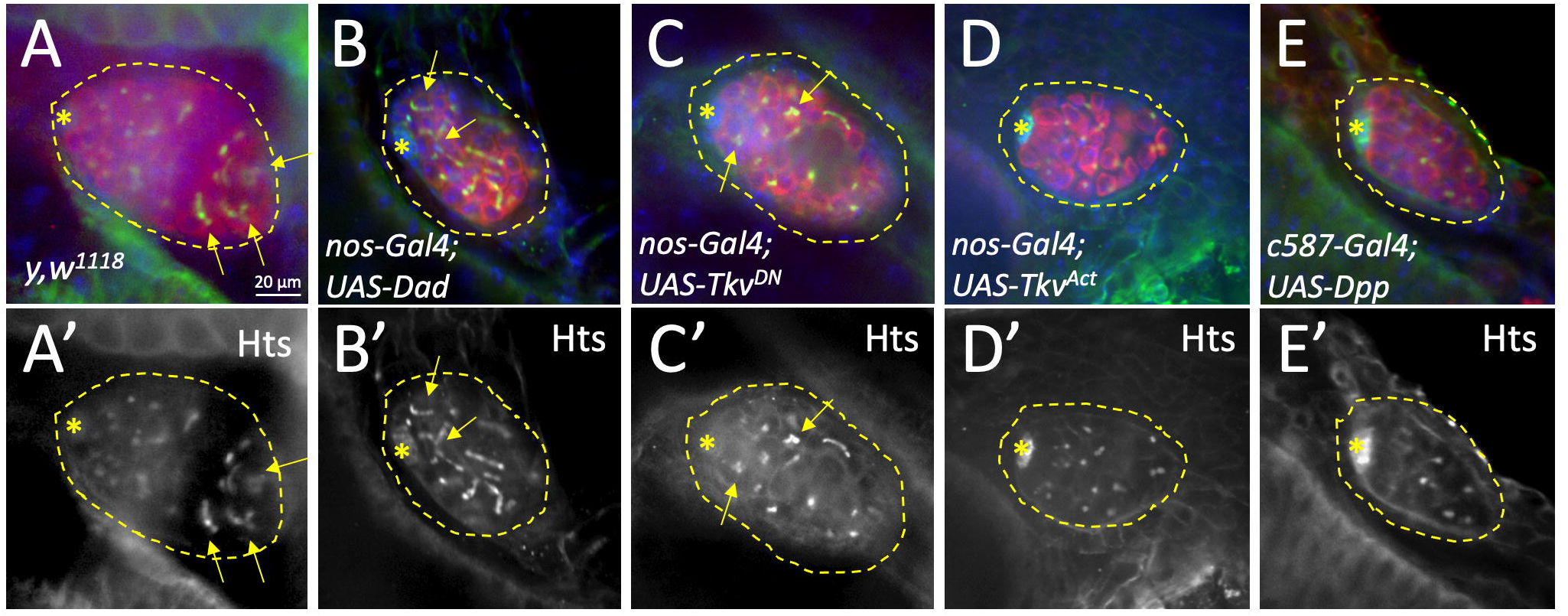
BMP signaling in the germline is necessary and sufficient for GSC self-renewal during early testis niche maturation. Immunostaining of late-L1/e-L2 testes with the fusome marker Hts (green or alone in grayscale; A’-E’) to detect germ cell differentiation status, as well as Vasa (red) and Fas3 (blue) to detect the germ cells and the hub, respectively. Testes outlined with asterisk indicating location of the hub. Scale bar is 20 µm. (A) y,w^1118^ control testis (n=31) showing normal fusomes with dot morphology in GSCs at the hub and branched fusomes in differentiating spermatogonia at the testes posterior. (B) nos-Gal4; UAS-Dad and (C) and nos- Gal4; UAS-Tkv^DN^ testes with BMP signaling inhibited specifically in the germline show elongated and branched fusomes in germ cells near the hub (yellow arrows; n=8, each), while (D) nos-Gal4; UAS-Tkv^Act^ and (E) c587-Gal4; UAS-Dpp testis with ectopically activated BMP lack germ cells with branched fusomes (n=19 and 15, respectively).

To determine whether BMP signaling is sufficient to promote GSC maintenance during early testis niche maturation, we next examined the impact of BMP hyperactivation on germline differentiation status (Figs 2D-E and S3). If BMP signaling is sufficient to promote GSC maintenance in larval testes, mis-activation of BMP in germ cells throughout the testes would be expected to prevent spermatogenic differentiation in the posterior of late-L1/early-L2 testes. We, therefore, assessed the impact of over-expressing a constitutively activated Tkv allele, Tkv^Act^ [70, 71], on fusome maturation. Similar experiments were also performed after ectopic expression of the BMP-activating ligand, Dpp, with the somatic gonadal *c587-Gal4* driver (refs from [57, 72]). Upon ectopic Tkv^Act^ expression in the germline, we found fusome branching to be absent in 79% of late-L1/early-L2 testes (Figs 2D and S3) while fusome maturation was ubiquitous in posterior germ cells within age-matched controls (Figs 2A and S3). Results were even more significant after somatic Dpp over-expression, with 93% of late-L1/early-L2 testes lacking branched or elongated fusomes (Figs 2E and S3). Thus, BMP activation in the germline is also sufficient to repress GSC differentiation in early stages of larval development. Together, these loss- and gain- of function studies directly show that BMP signaling in the germline plays a critical role in the maintenance and self-renewal of GSCs during early larval testis niche maturation.

### Germline repression of BMP signaling in late-stage larvae

While our analyses focus on BMP function in the germline during early stages of testes niche formation/maturation, we also assessed the impact of altered BMP signaling in later stages of testes development (S4 Fig). Interestingly, while ectopic expression of constitutively active Tkv in the germline inhibits spermatogonial differentiation in late L1/early L2 testes (Figs 2D and S4B), differentiation is recovered in L3 testes. Specifically, the majority of L3 testes (73%) resemble controls with a clear anterior to posterior progression of spermatogonial differentiation and branched fusomes at the testis posterior (S4C Fig). These observations are consistent with findings that the BMP feedback inhibitor, Smurf, downregulates BMP signaling in germ cells of 3^rd^ instar larvae and adult testes [61].

### BMP signaling is required in somatic gonadal cells for spermatogonial differentiation

Because pMad staining is detected in somatic cells of late-L1/early-L2 testes (Figs 1E and S2), we also assessed whether BMP signaling is required in somatic cells during early testes maturation (Figs 3 and S5). To do so, BMP signaling was inhibited in somatic cells of the testis, and the effect on late-L1/early-L2 larval testes examined. Upon ectopic expression of either Dad or Tkv^DN^ in testis somatic cells (Fig 3B-C), the testis anterior appears morphologically normal with GSCs containing dot fusomes arrayed around the hub and early signs of fusome elongation in adjacent germ cells. However, somatic BMP inhibition caused significant defects in spermatogonial differentiation, with germ cells at the testes posterior displaying spherical fusomes and/or reduced fusome branching in 100% and 82% of testes over-expressing Dad or Tkv^DN^, respectively (Figs 3B-C and S5). Interestingly, expression of the constitutively activated Tkv^Act^ allele in testes somatic cells also resulted in germ cell differentiation defects, with dot fusomes detected in the testes posterior in 90% of testes examined (Figs 3D and S5). Thus, consistent with detection of pMad staining in somatic cells of early-L2 testes (Figs 1 and S2; [61], not only is BMP signaling required in somatic cells for spermatogonial differentiation, but proper levels of BMP signaling in somatic cells appear to be critical for this process. Somatic BMP activation likely functions in a manner similar to the adult testes [73] where it promotes differentiation of cyst stem cell progeny as they are displaced from the testis niche.

**Fig 3.**
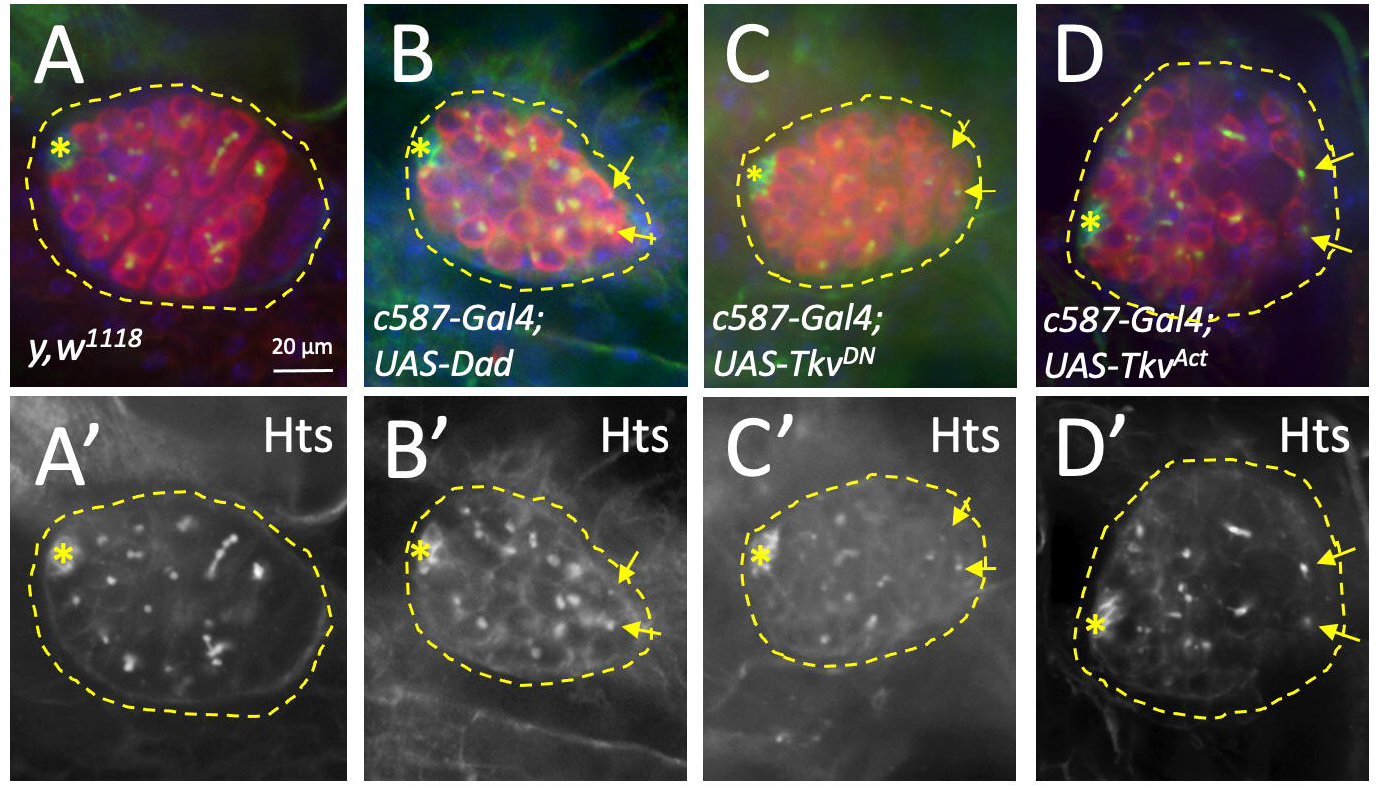
Somatic BMP signaling is required for normal spermatogenic differentiation. Immunostaining of late-L1/e-L2 testes with the fusome marker Hts (green or alone in grayscale; A’-D’) to detect germ cell differentiation status, as well as Vasa (red) and Fas3 (blue) to detect the germ cells and the hub, respectively. Testes outlined with asterisk indicating location of the hub. Scale bar is 20 µm. (A) y,w^1118^ control testis showing branched fusomes in the testis posterior consistent with spermatogenic differentiation (n=31). (B) c587-Gal4; UAS-Dad and (C) c587-Gal4; UAS-Tkv^DN^ testes with BMP signaling inhibited in the CySC lineage (n=24 and 11, respectively), as well as a (D) c587-Gal4; UAS- Tkv^Act^ testis with ectopic BMP activation (n=10), show lack of fusome branching and dot-shaped fusomes consistent with undifferentiated germ cells (arrows) in the testis posterior.

### CySCs activate BMP signaling in germ cells during testis niche maturation

We next assessed how BMP is activated in larval germ cells during early testis niche maturation. Attempts to detect BMP ligand expression in larval testes via *in situ* hybridization were unsuccessful (data not shown) so we used alternative methods. Previously, we have shown that hyperactivation of the Jak-STAT signaling pathway in somatic cells of larval testes causes expansion of CySCs throughout the testes coupled with reduced spermatogenic differentiation [57]. Consistent with this observation, ectopic expression of a constitutively activated Jak allele (Jak^Act^) in the CySC lineage results in somatic cells expressing the CySC marker Zinc Finger Homeodomain-1 (ZFH-1) throughout L2 testes, whereas ZFH-1 staining is restricted to CySCs docked at the hub in controls (Fig 4A-B). The spermatogenic differentiation defect observed after somatic Jak^Act^ over-expression is marked by loss of fusome branching and an expansion of under-differentiated germ cells with dot-fusomes (Fig 4A-B).

**Fig 4.**
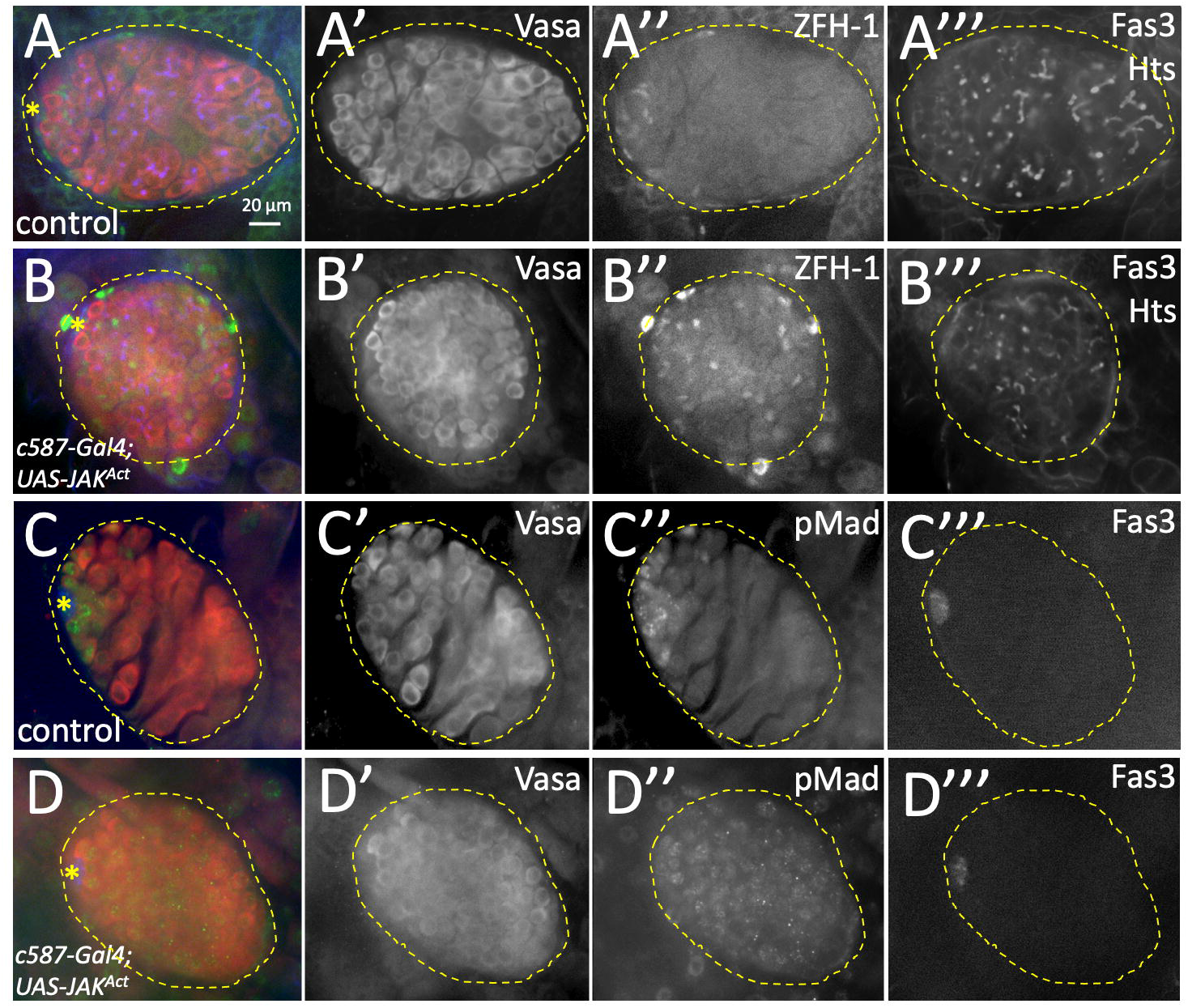
Germline BMP activation is controlled by signals from the CySCs. Immunostaining of e-L2 testes with either ZFH-1 (A,B; green) to detect CySCs or pMad (C,D; green) to detect BMP activation, as well as Vasa to detect germ cells (A-D; red), Fas3 to detect hub cells (A-D; blue), and Hts for fusomes (A,B; blue). Individual channels shown in gray scale. Testes outlined and scale bar is 20µm. (A) c587-Gal4/+ sibling control testis show normal fusome morphology and ZFH-1 expression restricted to CySCs adjacent to the hub (n=24). (B) c587-Gal4/+;UAS-Jak^Act^/+ testis with undifferentiated germ cells that lack branched fusomes as well as expansion of ZFH-1 positive somatic CySCs throughout the testes (n=23). (C) c587-Gal4/+ sibling control testis with pMad restricted to GSCs adjacent to the hub (n=12). (D) c587- Gal4/+; UAS-Jak^Act^/+ testis with pMad expression in germ cells throughout the testis (n=10).

As BMP signaling from CySCs to germ cells promotes GSC self-renewal in the adult testes, we therefore utilized larval testes showing CySC expansion due to somatic Jak^Act^ mis- expression to assess if CySCs are a source of BMP activating ligands during larval testes maturation. Specifically, we examined the ability of Jak hyperactivation to alter the pattern of pMad staining in the germline of L2 testes. We find that ectopic Jak^Act^ expression in the CySC lineage results in pMad staining in germ cells throughout the testes, while pMad is restricted to anterior germ cells in sibling controls (Fig 4C-D). Notably, hub cells, which have also been shown to be a source of BMP ligand secretion in the adult testes, remained restricted to the testes anterior after Jak hyperactivation (Fig 4C-D). Together with observations presented above that (i) BMP signaling is necessary and sufficient for GSC maintenance during early testis niche maturation and (ii) Jak hyperactivation causes expansion of CySCs and under-differentiated germ cells, these data indicate that BMP signaling from the CySCs to germ cells plays a critical role in controlling GSC self-renewal in the newly formed larval testis niche. While this mechanism is similar to that in adult testes, phenotypic recovery in L3 testes after germline BMP hyper-activation (Fig 3C) and Smurf-mediated downregulation of BMP activation in L3 and adult testes [61] show that BMP signaling from CySC to the germline has a more prominent role in regulation of GSC maintenance during early testis niche maturation than at later stages of developmental or in mature adults.

### BMP signaling is dynamically activated and required for fusome maturation during GSC stem cell niche formation

Finally, we sought to examine the role of BMP signaling in GSCs during testis niche morphogenesis. To do so, we first characterized BMP activation in late-stage embryonic gonads. Specifically, stage 16 through early-L1 testes were immunostained to detect the pattern of nuclear pMad in the germline (Figs 5A-Eand S1). At stage 16 (Fig 5A, ∼15 hrs AEL), pMad is detected at low levels in the cytoplasm of germ cells. This observation is consistent with prior work examining pMad staining during earlier stages of gonad formation [27]. As GSC niche morphogenesis progresses, nuclear pMad is detectable with a dynamic staining pattern. In early stage-17 testes (15-18 hrs AEL), nuclear pMad is initially detected in germ cells at the testis posterior (Figs 5B and S1). Subsequently, nuclear pMad staining extends anteriorly in mid/late-stage-17 testes (Figs 5C and S1; 18-21 hrs AEL), followed by a broader staining pattern with nuclear pMad in germ cells throughout late-stage 17/early-L1 testes at the embryo to larval transition (Fig 5D, 21-24 hrs AEL). Once a functional GSC niche has formed in early-L1 larvae (24-28 hrs AEL), the pattern of nuclear pMad becomes restricted to anterior germ cells as previously discussed (Figs 1C-D, S1, and 5E). While not all testes at similar stages of embryogenesis show the exact same pattern of nuclear pMad expression, the trend of posterior pMad, followed by an anterior shift, and subsequent restriction to the newly formed testes niche is observed in the majority of testes (S1 Fig).

**Fig 5.**
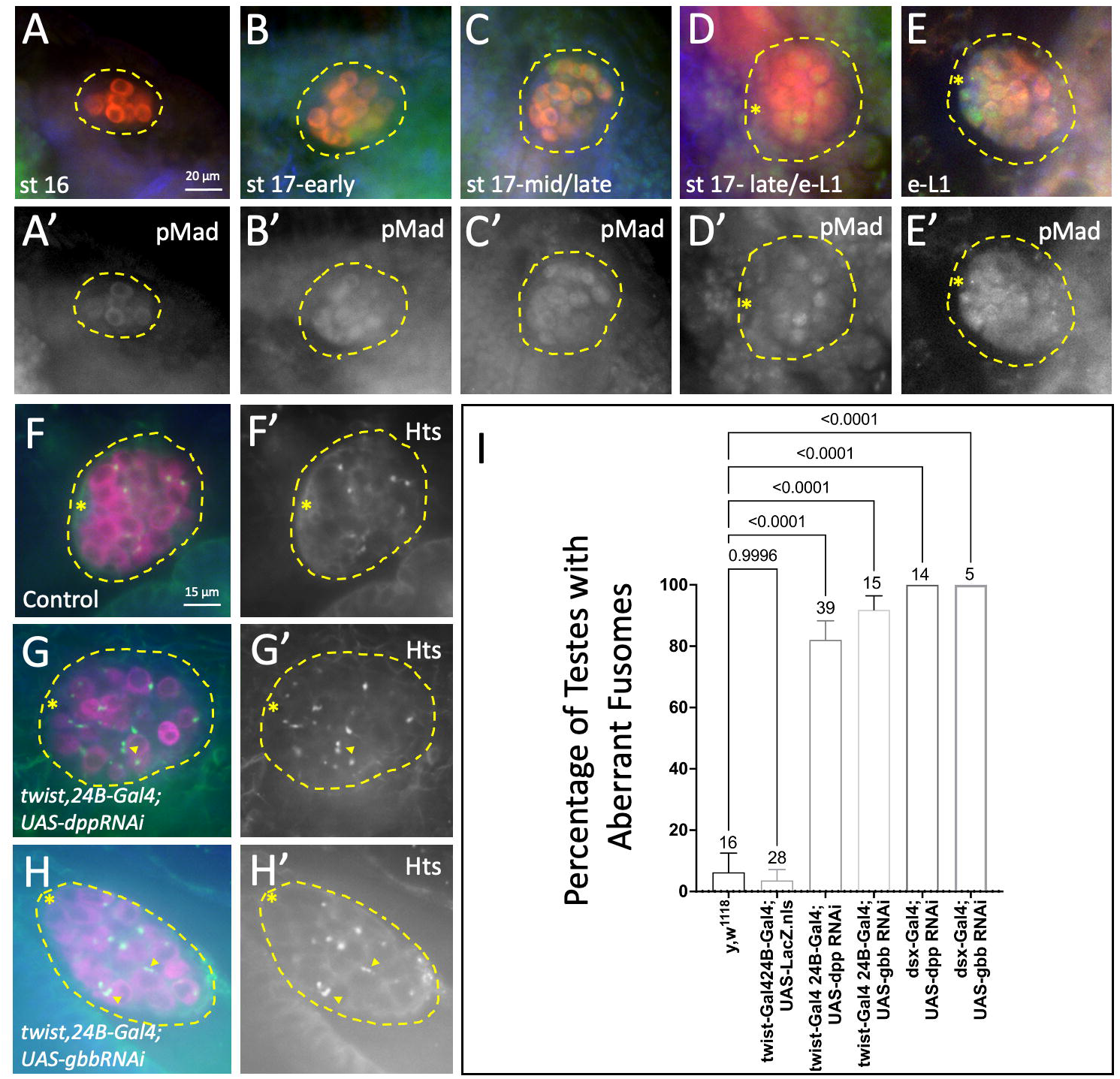
BMP is dynamically activated in germ cells during testis niche morphogenesis. (A-E) *Dynamics of BMP activation* - Immunostaining of wild type embryos and e-L1 larvae 12- 28 hrs AEL for BMP activity as assessed by pMad (green; A’-E’ alone in grayscale) as well as Vasa (red) and Fas3 (blue) to detect germ cells and the hub cells, respectively. (A) stage 16 embryonic gonad lacking nuclear pMad in the germline (∼15 hrs; n=8). (B) early-stage 17 testis with nuclear pMad in a posterior germ cells (15-18 hrs; n=13). (C) mid/late -stage 17 with nuclear pMAD in posterior germ cells extending anteriorly (18-21 hrs; n=20). (D) late-stage 17/early-L1 testis at the embryo to larval transition with nuclear pMad in a subset of germ cells throughout the gonad (21-24 hrs; n=15). (E) e-L1 testis with nuclear pMad restricted to the newly established GSC niche (24-28 hrs AEL; n=16). (F-I) *Impact of somatic BMP knockdown -* Immunostaining of e-L1 larvae (22-28 hrs AEL) with the fusome marker Hts (F-H, green; F’-H’ in grayscale) to detect fusome morphology, as well as Vasa (red) and Fas3 (blue) to detect germ cells and the hub, respectively. (F) Control y,w^1118^ testis with germ cells containing dot-shaped fusomes (n=16). (G) twist-Gal4, 24B-Gal4; UAS-dpp-RNAi and (H) twist-Gal4, 24B-Gal4; UAS-gbb-RNAi testes with individual germ cells showing aberrant fusomes with fragmented and/or elongated morphology (arrow heads; n=39 and 15, respectively). (I) Average percentage of testes in e-L1 larvae showing aberrant fusomes in controls and in samples with somatic knockdown of BMP-activating ligands. Standard error, sample size, and p-values from ANOVA with Tukey post-hoc comparisons shown. Testis outlined in all images, with asterisk indicating location of the hub. Scale bar at 20 or 15 µm (A-E or F-H, respectively).

### BMP signaling is required for fusome maturation during testis niche morphogenesis

To better understand the role of BMP activation during testis niche morphogenesis, we also assessed the impact of altered BMP signaling on germ cell behavior and niche formation. Because loss of BMP signaling in the germline was previously shown to impact PGC migration [27], experiments were designed to inhibit germline BMP activity after germline-SGP contact. Specifically, the somatic *twist-Gal4, 24B*-Gal4 driver combination was used to express UAS-RNAi constructs against the BMP activating ligands, Dpp or Gbb, in embryonic SGPs [74, 75]. Late stage 17 through early-L1 testes (21-28 hrs AEL) were then immunostained to assess for changes in gonad as well as fusome morphology (Fig 5F-I). We also examined Mad staining to confirm that RNAi against Dpp or Gbb inhibited BMP signaling. Interestingly, somatic RNAi against either Dpp or Gbb resulted in the absence of nuclear pMad in germ cells within the majority of testes (Figs S6-7). Further, while Dpp or Gbb knockdown does not alter the characteristic rosette morphology of the early larval testes niche, we find that somatic knockdown does cause defects in fusome morphology within the newly formed niche (Fig 5F-I). Specifically, somatic Dpp or Gbb knockdown frequently resulted in germ cells with mis-shapen and/or fragmented fusomes, while such defects were rare in controls. Indeed, 82% of testes showed fusome defects after Dpp knockdown induced by *twist,24B-Gal4*, and aberrant fusomes were observed in 92% of testes after Gbb RNAi knockdown (Fig 5F-I). Similar results were also observed after Dpp knockdown with the SGP-specific Gal4 driver, *dsx-Gal4* (Fig 5I). Conversely, aberrant fusomes were observed in only 6% and 7% of control testes (wild type or *twist,24Bgal4;UAS-LacZ*, respectively; Fig 5F-I). While not all germ cells within Dpp or Gbb knockdown testes showed mis-shapen or fragmented fusomes, the increase was statistically significant compared to controls (S8 Fig). Notably, fusome defects were observed in single germ cells and did not appear to span the cytoplasm of interconnected germ cells; indicating defects in fusome maturation rather than the early onset of spermatogonial differentiation due to absence of BMP activity. Taken together, these data indicate that BMP signaling from SGPs to the germline in late embryonic testes serves to promote normal maturation of fusomes that are an essential aspect of germ cell identity and spermatogonial differentiation during testis niche development.

## Discussion

In this study, we directly examined the function and dynamics of BMP signaling in *Drosophila* testes during embryonic GSC niche morphogenesis as well as testis maturation in early larvae. During early stages of testis niche maturation, we find that germline BMP activity is restricted to GSCs and early gonialblast cells. We also provide direct evidence that BMP signaling in the germline is both necessary and sufficient for maintenance of a self-renewing GSC population. Furthermore, our evidence indicates that newly formed CySCs are a source of BMP activating ligand during early niche maturation. As development progresses, however, BMP activation is no longer sufficient to alter GSC self-renewal. Additionally, we find that BMP signaling is activated in differentiating somatic cyst cells during early niche maturation. This BMP function is critical for early somatic cyst cell function as evidenced by defects in spermatogonial differentiation in larvae when BMP activity is altered in somatic cells. Finally, we find that BMP signaling shows a dynamic pattern of activation in the germline during embryonic niche morphogenesis. Loss of BMP signaling from the soma to the germline impacts germ cell maturation as indicated by defects in fusome morphology. These data confirm and build upon prior studies indicating that BMP signaling is critical for promoting germ cell identity in PGCs during embryonic gonad formation [27], as well as observations that Smurf downregulates BMP signaling as testis niche maturation progresses [61].

Together, these data provide direct evidence of a key role for BMP signaling in the newly formed GSC niche that is similar in function to the adult testis niche. In both systems, BMP ligand secretion from CySCs to adjacent GSCs prevents spermatogonial differentiation and helps to maintain the GSC population within the niche [15,46,76,77]. Jak-STAT signaling from the hub to the GSCs and CySCs also helps to maintain these stem cell populations in the adult and larval niche [45,56,57,78,79]. Similarities between the two systems indicate that the larval niche can provide a highly tractable system to elucidate additional mechanisms controlling testis stem cell maintenance and differentiation that might not be identified in the adult due to lethality.

Our observation that BMP signaling is required in developing somatic cyst cells is also consistent with BMP function in adult cyst cells. However, while the mode of BMP action in adult GSCs is well understood [80, 81] less is known about BMP function in cyst cells. It is known that BMP signaling is activated non-autonomously in adult cyst cells, and it has been proposed that activation triggers signaling from cyst cells to the adjacent germline to restrict proliferation of differentiating germ cells [73,80,81]. BMP activity in adult cyst cells is also mediated by the type I and II BMP receptors, Sax and Punt, as well as the transcription factors, Schnurri (Shn) and dSmad2/Smox [73, 82]. However, the ligand activating BMP signaling in somatic cyst cells has not been identified. Since dSmad2/Smox typically mediates TGF-β signaling through the Activin rather than BMP pathway, BMP signaling in cyst cells may not proceed through standard mechanisms. Additionally, the BMP receptor, Tkv, has been proposed to serve as a ligand sink that prevents BMP activation in germ cells localized away from the niche so that they can initiate spermatogonial differentiation [64]. Additional studies in the adult and larval testis stem cell niches are, therefore, necessary for us to better understand BMP function in somatic cyst cells critical for fertility.

Since the adult and newly formed testis niches function similarly, it is intriguing that the effects of BMP signaling in 3^rd^-instar larval and adult testes are less robust than in the newly formed testis niche. While it is possible that the difference in our study results from reduced activity of the *nanos-Gal4* driver as development progresses, this observation is also consistent with the downregulation of BMP signaling in the germline by the E3-ligase, Smurf [61]. So, why would BMP play a greater role during early development? One hypothesis is that germline differentiation must be downregulated in the early niche to allow time for formation and morphogenesis of cell types in the testes posterior that are necessary for later stages of spermatogenesis. Indeed, while the testis niche becomes functional at the embryo to larval transition, and cyst cell as well as spermatogonial differentiation is observed in mid-1^st^ instar larval testes [54,56,57], the testes posterior remains relatively undifferentiated even in 3^rd^ instar larval testes. Furthermore, connections are not made to other structures in the reproductive tract formed from the genital disc until early pupal development [83, 84]. Thus, downregulation of BMP signaling in 3^rd^ instar larvae may correlate with early stages of somatic cell differentiation in the testis posterior that are required for spermatogenic progression. Further studies examining development of somatic cells in the testis posterior are necessary to validate this hypothesis. Together, these analyses will shed light on the interplay between signaling networks that control differentiation and morphogenesis of the *Drosophila* testis that likely have implications for mammalian organ systems.

Additional work must also be performed to understand how and why BMP signaling is activated in posterior germ cells at the end of embryogenesis just prior to formation of a functional testis stem cell niche. Our observation that loss of BMP signaling during embryonic niche morphogenesis results in defects in fusome morphology is consistent with prior work indicating that BMP promotes germ cell identity required for fusome formation during gonad coalescence [27]. However, defects we observe in our study do not appear to be biased toward the testis posterior (data not shown). Thus, it is possible that BMP activation in posterior germ cells plays another role that we have not yet uncovered. Given the importance of BMP signaling in GSC maintenance, and the fact that cyst cell differentiation is not observed in the testis until mid-larval-1^st^ instar, one possibility is that posterior BMP activation serves to repress germ cell differentiation that would result in the formation of germ cell tumors in the absence of functional cyst cells. While we did not observe premature spermatogenic differentiation after RNAi-inhibition of either the Dpp or Gbb ligands in our study, it is possible that the simultaneous loss of both these ligands may elicit these effects. It will also be interesting to ascertain which cells in the testis are responsible for BMP activation in the testes posterior, as well as how the positioning and morphology of these cells result in the observed dynamic pattern of BMP activity in germ cells during testis stem cell niche morphogenesis in late embryos.

In summary, we directly show a role for BMP signaling in GSC maintenance and cyst cell function during early testis niche maturation. Our work contributes to the broader understanding of mechanisms by which cell signaling controls the dynamic process of testis stem cell niche development. Additionally, defects in fusome morphology observed after BMP inhibition during niche morphogenesis highlight the importance of BMP signaling in promoting germ cell fate. The dynamic pattern of BMP activity in germ cells in late stage embryos points toward additional functions for BMP signaling prior to niche formation. Going forward, similarities between the adult and developing testes suggest that testes development can provide a tractable system for elucidating mechanisms controlling stem cell behavior that might be more difficult to identify in adults. At the same time, differences in mechanisms controlling testis development can provide information as to how stem cell systems must be adapted to not only maintain stem cell types, but allow for tissue growth and integration into surrounding tissues. Therefore, further analyses of *Drosophila* testes development will likely yield information with broader implications for mammalian organ development and reproduction.

## Materials and methods

### Fly stocks

y, w^1118^ flies were used as controls unless otherwise noted. nos-Gal4::VP16 [85] was employed to induce germline expression of UAS-transgenes, while the c587-Gal4 [72], dsx-Gal4 [86] and twist, 24B-Gal4 drivers [74, 87] were used to express UAS-transgenes in the soma. UAS-constructs used to modulate BMP activity include: UAS-Dpp [88]; BDSC stock 1486), UAS-Dad [68], UAS-Tkv^Activated (Act)^ [70, 71], UAS-Tkv^DN^ [69], UAS-dpp-RNAi [89]; BDSC stock 33618/TRiP.HMS00011), UAS-gbb-RNAi [89]; BDSC stock 34898/TRiP.HMSS01243). Other UAS-constructs include: UAS-LacZ::GFP.nls (BDSC stock 6451), and UAS-Jak^Act^ to over-express a constitutively activated version of JAK, Jak^Act^ [22, 90]. For all matings, virgin females were collected from flies carrying the Gal4 driver. All fly stocks were obtained from the Bloomington Stock Center (http://flystocks.bio.indiana.edu/) unless otherwise specified.

### Sample collection

Embryos and larvae were collected on apple juice plates and fixed according to Fidler et al (2013). Samples were collected at the following time increments: 0-22 hrs AEL (embryos), 22-28 hrs AEL (late stage 17 embryos/early L1 larvae), 22-48 hrs AEL (L1 larvae), 48-72 hrs AEL (L2 larvae), 72-96 hrs AEL (L3 larvae) at 24°C, with embryo staging performed according to [91]. Alternatively, the following time increments were used for fine-scale analysis of pMad expression: 0-18 hrs AEL (stage 1 – early stage 17 embryos), 18-19 hrs AEL (mid stage 17 embryos), 19-21 hrs AEL (late stage 17 embryos), 21-24 hrs AEL (late-stage 17 embryos/early L1 larvae), 24-32 hours AEL (early/mid-L1 larvae), 32-40 hours AEL (mid-L1 larvae), 40-48 hours AEL (late-L1 larvae), 48-60 hrs AEL (early-L2 larvae). To collect samples with Jak activity induced in the soma, samples were collected with a temperature upshift as follows: Embryos collected on apple juice plates from 0-7 hrs AEL at 24°C and then incubated at 29°C for 48-60 hrs (early-L2). Genotyping of c587-Gal4/+; UAS-Jak^Act^/+ versus c587-Gal4/+; +/+ sibling controls from these collections was determine either by expansion of ZFH-1 staining in somatic cells or by presence of under-differentiated germ cells with spherical morphology throughout the testes as previously reported [57]. In all instances, virgin females carrying the Gal4 driver were mated to males carrying the UAS- transgene.

### Antibodies and immunostaining

Immunostaining of embryos and larvae was performed as described [92]. The following primary antibodies were used to detect germ cells: chick anti-Vasa at 1:3000 (K. Howard), rabbit anti-Vasa at 1:5000 (R. Lehmann) and rat anti-Vasa at 1:40 (Developmental Studies Hybridoma Bank [DSHB]). BMP activation was detected with rabbit anti-pMad at 1:100 (Epitomics; #1880/EP823Y). Additionally, mouse anti-1B1/Hts at 1:4 (H. Lipshitz; DSHB) was used to detect germ cell differentiation status, and hub cells were visualized using mouse anti-Fasciclin3 (FasIII) at 1:10 (C. Goodman; DSHB) or rat anti-DN-Cadherin (DN-Ex #8) at 1:20 (T. Uemora; DSHB). Cyst stem cells were detected with rabbit anti-ZFH-1 at 1:5000 (R. Lehman). The following secondary antibodies (Invitrogen) were used: goat anti-chick 546, goat anti-rabbit 546, goat anti-rabbit 488, goat anti-rat 546, goat anti-rat 488, goat anti-mouse 633 and goat anti-mouse 488. All secondary antibodies were used at a 1:500 final dilution. To enhance staining of pMad and ZFH-1, samples were incubated with rabbit anti-pMad or anti-ZFH-1 alone for ∼24 hours at 4°C, and then an additional 4 hours at room temperature. Samples were then rinsed, washed, and goat anti-rabbit fluorescent secondary antibody added overnight at 4°C. Samples were once again rinsed, washed, and blocked, and then subjected to immunostaining with other antibodies in the absence of anti-pMad or ZFH-1as per usual [92] with goat anti-rabbit secondary against antibody added with other secondary antibodies. To stain for nuclei, DAPI (Roche) was used at 1:1000 after completion of final secondary antibody washes.

### Microscopy

Samples were mounted in 70% glycerol containing 2.5% DABCO (Sigma) and *p-* phenylenediamine anti-fade agent (Sigma) at a final concentration of 0.2 mg/mL. Slides were viewed with an Olympus BX51 microscope equipped with a DSU spinning disc confocal system and Q-imaging RETIGA-SRV CCD camera. Images were captured and analyzed with Slidebook software by 3I.

### Statistical analyses

To assess for changes in BMP activity in the developing germline, wild type samples of different ages were scored for presence of nuclear pMad as follows: Posterior pMad (1 or more germ cells with nuclear pMad in posterior half of the testis but no stain above background elsewhere), extending anterior (presence of nuclear pMad in germ cells located in the posterior half of the testis as well as nuclear pMad staining in a subset of germ cells in the testis anterior), most germ cells (presence of strong nuclear pMad staining in germ cells throughout the gonad), anterior restricting/restricted (absence of nuclear pMad staining in germ cells at the testes posterior coupled with nuclear staining in the anterior). Alternatively, to assess for inhibition of BMP activation after BMP ligand knockdown, testes were scored for presence/absence of nuclear pMad in germ cells above background. Testes lacking nuclear pMad in all germ cell were scored as pMad negative, while those with 1 or more germ cells with nuclear pMad were scored as positive. In all instances, statistical significance of results was determined by chi square test with a p-value based on 2000 replicates of a Monte Carlo simulation.

To assess defects in fusome morphology, the number of testes with normal fusomes (dot fusomes in GSCs at the testis anterior and branched fusomes in the posterior) or testes with aberrant fusomes (dot fusomes in posterior or elongated/branched fusomes in testis anterior) were counted and the percentage of all testes was determined. The differences in the proportions of testes with each fusome morphology were assessed using chi square tests with a p-value based on 2000 replicates of a Monte Carlo simulation. The average proportion of testes with aberrant fusomes was assessed using analysis of variance (ANOVA).

To assess for changes in gonad size, the length of each testis oriented longitudinally within the imaging plane was measured using Slidebook 5.0 and compared to a set of control testes of comparable age. The significance of results was determined with an unpooled-variance two-tailed Student’s t-test. For all statistics, significance was assigned at or below p = 0.05.

## Supporting information

**S1 Fig. BMP activity is dynamic during early testis GSC niche development.** Heat map depicting changes in nuclear pMad distribution in germ cells of developing testis ranging from 12-60 hrs AEL (stage 15 embryos through early-L2). Percentage of germ cells with germline pMad distribution at different ages is indicated by degree of shading for the following categories: Posterior germ cells, extending anterior, most germ cells, anterior germ cells restricting/restricted as described in methods. Stage 5-16 embryos (12-15 hrs; n=15), stage 17-early (15-18 hrs; n=13), stage 17-mid/late (18-21 hrs; n=20), stage 17-late/e-L1 (21-24 hrs; n=15), early-L1 (24- 28 hrs; n=14), early/mid-L1(28-32 hrs, n=16), mid-L1 (32-40 hrs; n=19), mid-L1/early-L2 (40- 60 hrs; n=23).

**S2 Fig. BMP activity is detected in newly differentiated cyst cells during testis maturation.** Immunostaining of wild type e-L2 testes for BMP activity (n=11) as assessed by pMad (green or alone in grayscale; A’) co-stained with Vasa (red) and Fas3 (blue) to detect the germ cells and the hub, respectively. Testis outlined with asterisk indicating location of the hub. Yellow arrows indicate location of pMad-positive somatic cyst cells.

**S3 Fig. Impact of altered BMP signaling in germ cells on fusome morphology during testis niche maturation.** Stacked bar chart showing percent of testes from late-L1/e-L2 samples with normal versus defective fusomes consistent with lack of GSC maintenance (elongated or branched fusomes in anterior) or defects in germline differentiation (dot fusomes in posterior) as indicated in key (right). Genotypes on Y-axis, and results from chi-square analysis (see methods) on top right. Sample size shown with Figure 2.

**S4 Fig. BMP is no longer sufficient to repress spermatogenic differentiation during later testis niche maturation.** Immunostaining of testes with the fusome marker Hts (green or alone in grayscale; A’-C’) to detect germ cell differentiation status, as well as Vasa (red) and Fas3 (blue) to detect the germ cells and the hub, respectively. Testes outlined with asterisk indicating location of the hub. Scale bar is 20 µm. (A) *y,w^1118^* control testis with normal fusome morphology (n=31). (B,C) nos-Gal4;UAS-Tkv^Act^ testes with ectopic BMP activation in the germline lack branched fusomes at (B) late-L1/e-L2, but show branched fusomes throughout the testis posterior consistent with normal spermatogenic differentiation in (C) L3 testes (n=31 and 11, respectively).

**S5 Fig. Impact of altered BMP signaling in somatic cells on fusome morphology during testis niche maturation.** Stacked bar chart showing percent of testes from late-L1/e-L2 samples with normal versus defective fusomes consistent defects in germline differentiation (dot fusomes in posterior) as indicated in key (right). Genotypes on Y-axis, and results from chi-square analysis (see methods) on top right. Sample size shown with Figure 3.

**S6 Fig. BMP activation is inhibited by somatic knockdown of BMP ligands.** Immunostaining of late stage17/e-L1 samples (22-28 hrs AEL) for BMP activity as assessed by pMad (green; A’- E’ alone in grayscale) as well as Vasa (red) and Fas3 (blue) to detect germ cells and the hub cells, respectively. Testis outlined with asterisk indicating location of the hub. Scale bar is 20 µm. (A) twist-Gal4,24B-Gal4;UAS-LacZ.nls and (B) dsx-Gal4;UAS-LacZ.nls control testes show nuclear pMad staining in germ cells as expected (n=13 and 15, respectively), while nuclear pMad is undetectable in (C) twist-Gal4, 24B-Gal4;UAS-dpp-RNAi (n=38), (D) dsx-Gal4;UAS-dpp-RNAi (n=6), (E) twist-Gal4, 24B-Gal4;UAS-gbb-RNAi (n=18), or (F) dsx-Gal4;UAS-gbb-RNAi (n=13) testes where expression of BMP activating ligands are inhibited in developing somatic cells.

**S7 Fig. Quantitative analysis of BMP inhibition after somatic knockdown of BMP activating ligands.** Stacked bar chart showing percent of testes from late-st17/e-L1 samples with presence of absence of nuclear pMad in the germline as indicated in key (right). Genotypes on Y-axis, and results from chi-square analysis (see methods) on top right. Sample size shown with Figure 6.

**S8 Fig. Percentage of germ cells with aberrant fusomes after somatic BMP ligand knockdown.** Bar graph showing mean percentage of germ cells per late-stage 17/e-L1 testis (22- 28 hrs AEL) with aberrant fusomes (Y-axis) for each genotype (X-axis). Standard error, sample size, and p-value from ANOVA analyses showing statistically significant differences in controls versus testes with either Dpp or Gbb knocked down in somatic gonadal cells.

## Acknowledgements

We are grateful to all our colleagues who have supported our lab with antibodies, stocks and technical assistance. We would also like to acknowledge the Bloomington *Drosophila* Stock Center at Indiana University for maintaining and providing fly stocks, and the Developmental Studies Hybridoma Bank developed under the auspices of the NICHD and maintained by The University of Iowa, Department of Biology. We specifically thank members of the Wawersik lab for critical comments on this manuscript.

