## Supplementary figures and images for "Bone Morphogenetic Protein (BMP) signaling regulates germline stem cell self-renewal in the newly formed *Drosophila* testis stem cell niche"

### Supplemental Fig1

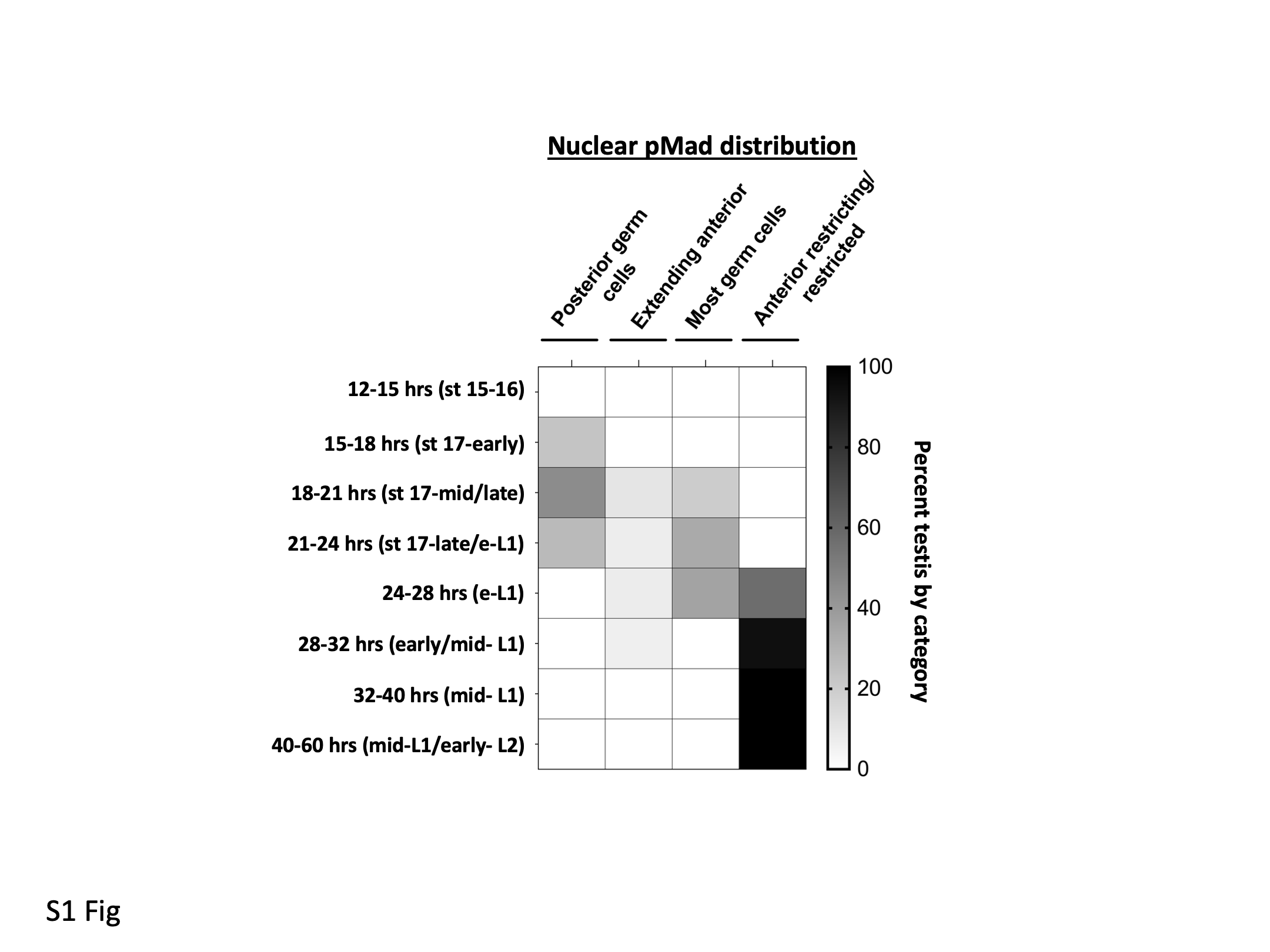

### Supplemental Fig2

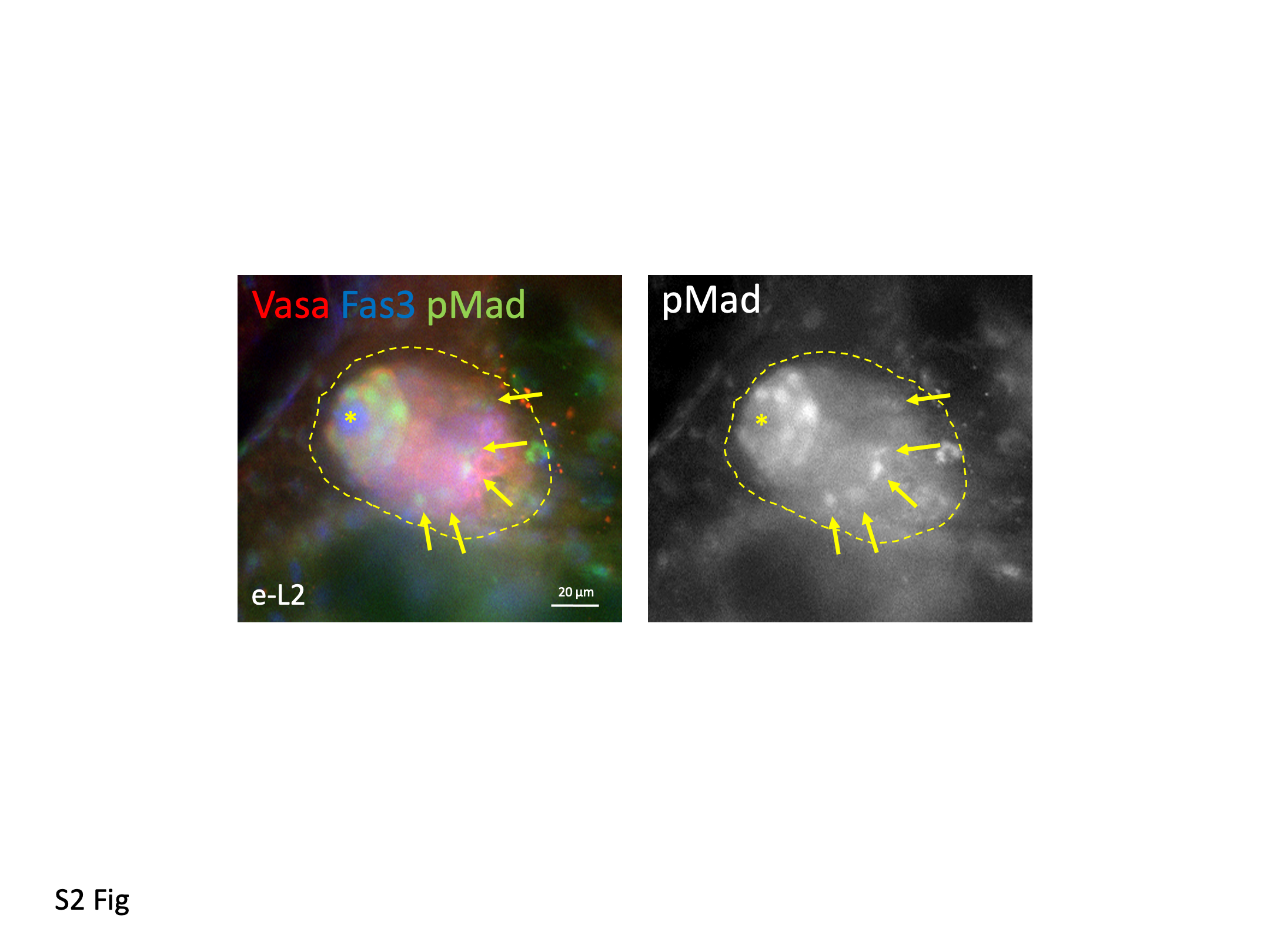

### Supplemental Fig3

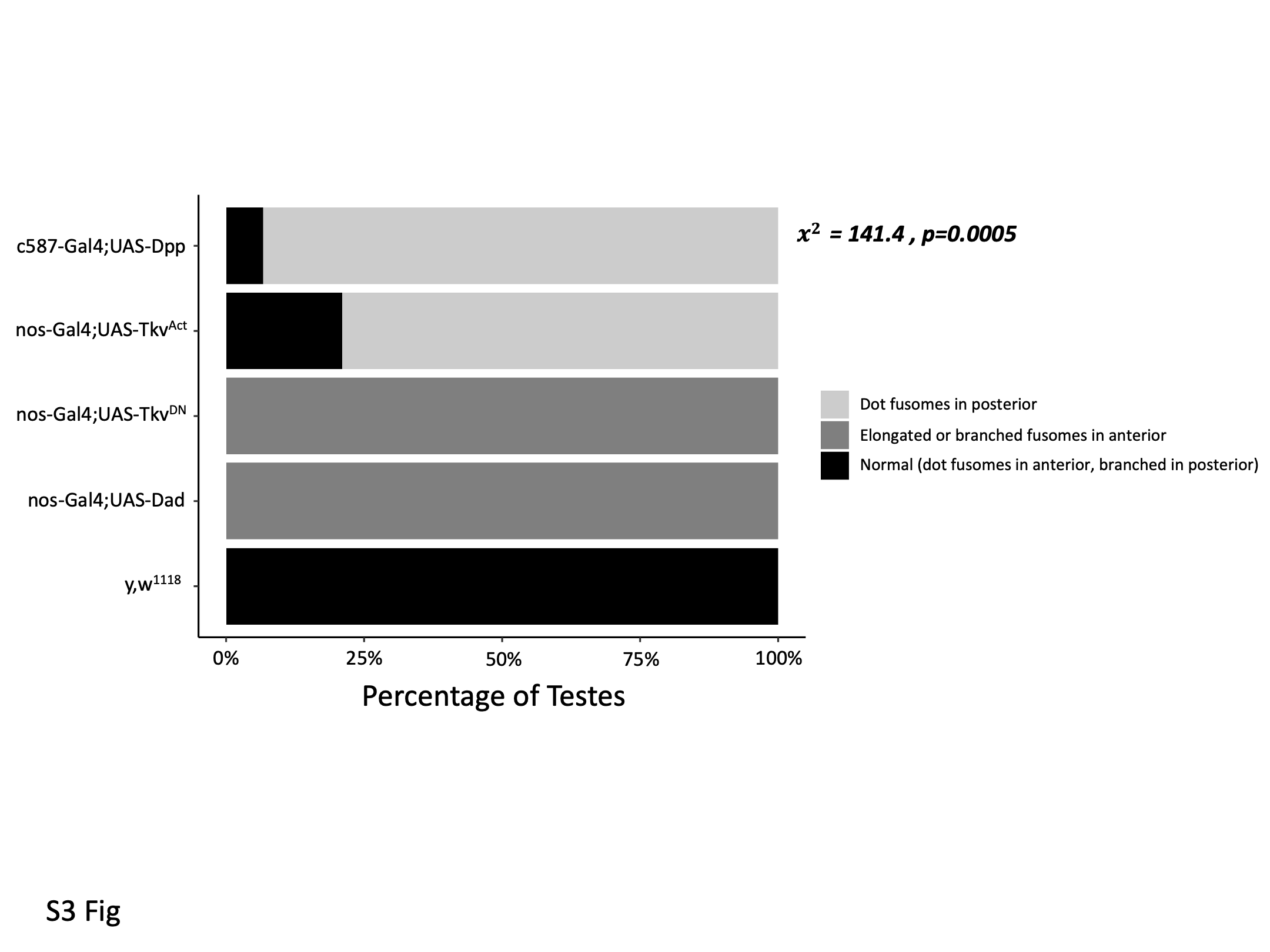

### Supplemental Fig4

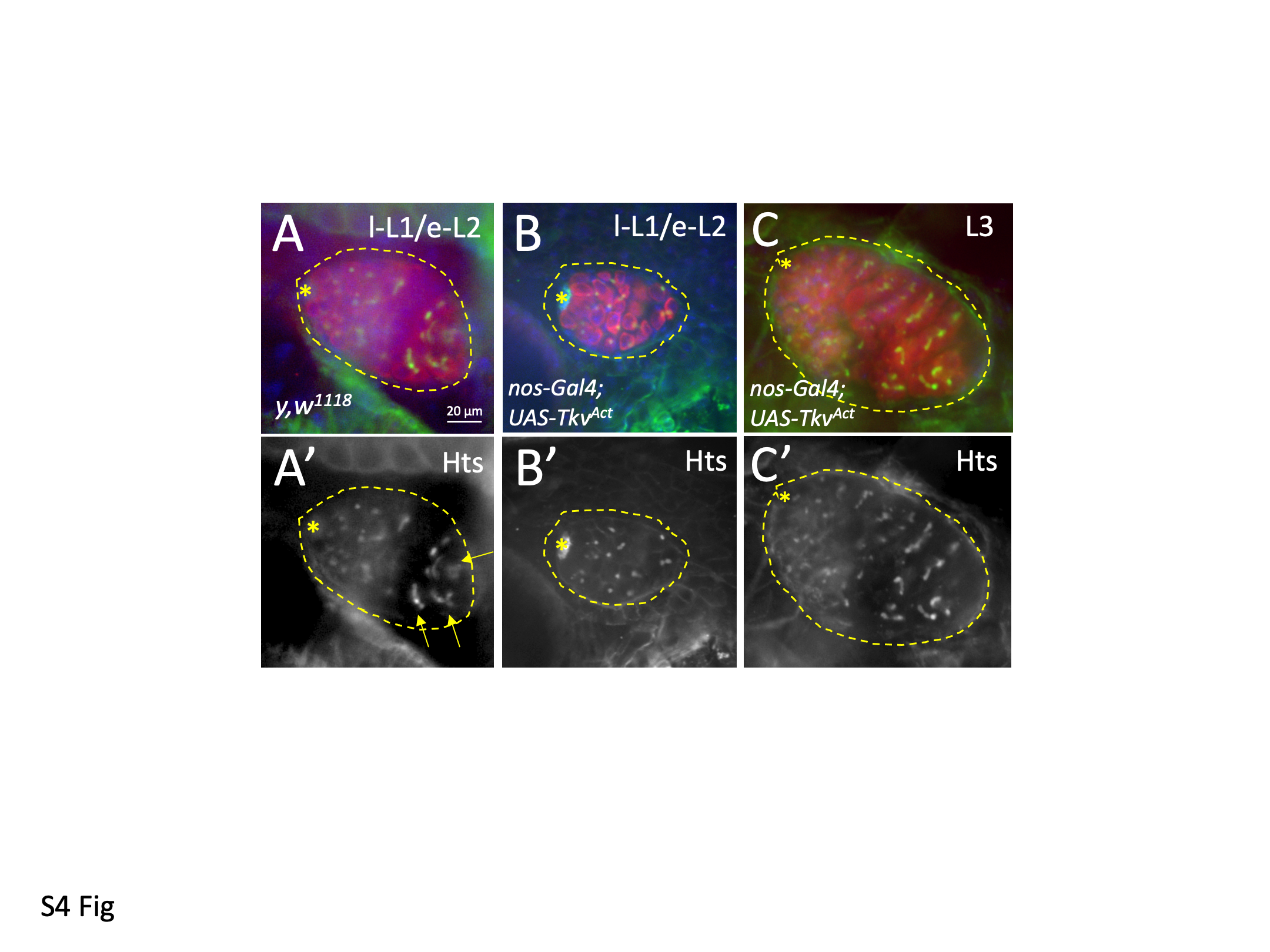

### Supplemental Fig5

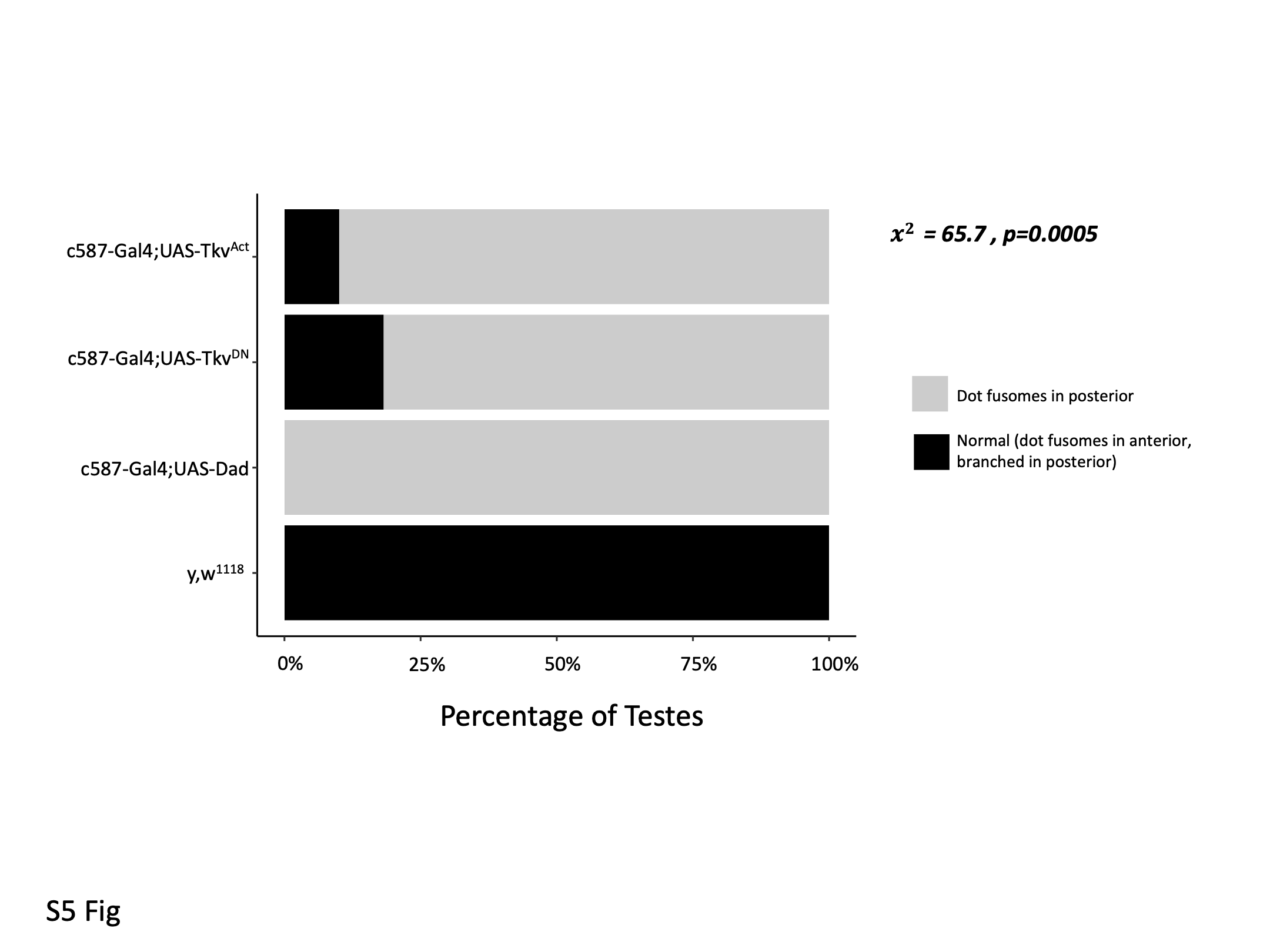

### Supplemental Fig6

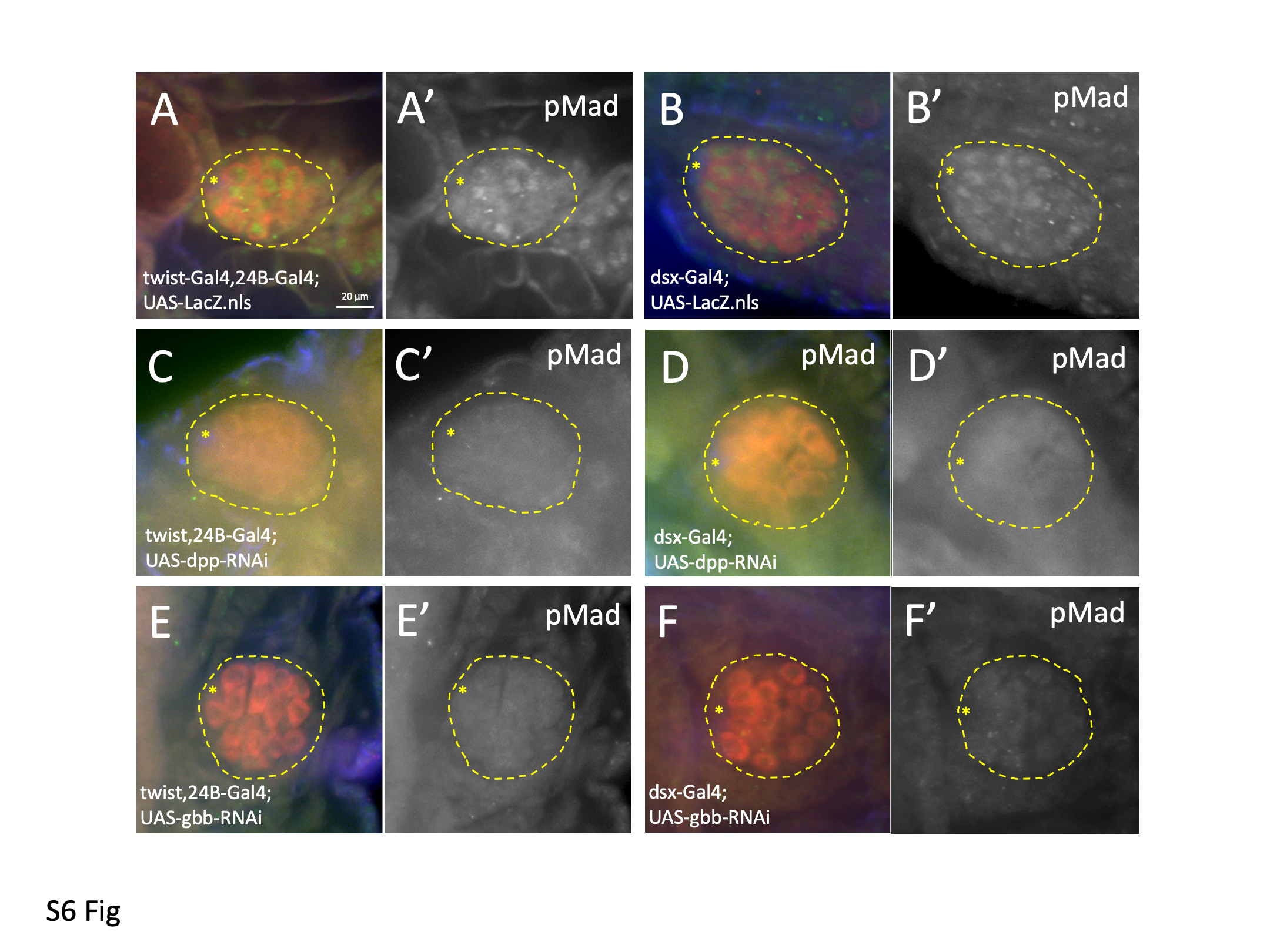

### Supplemental Fig7

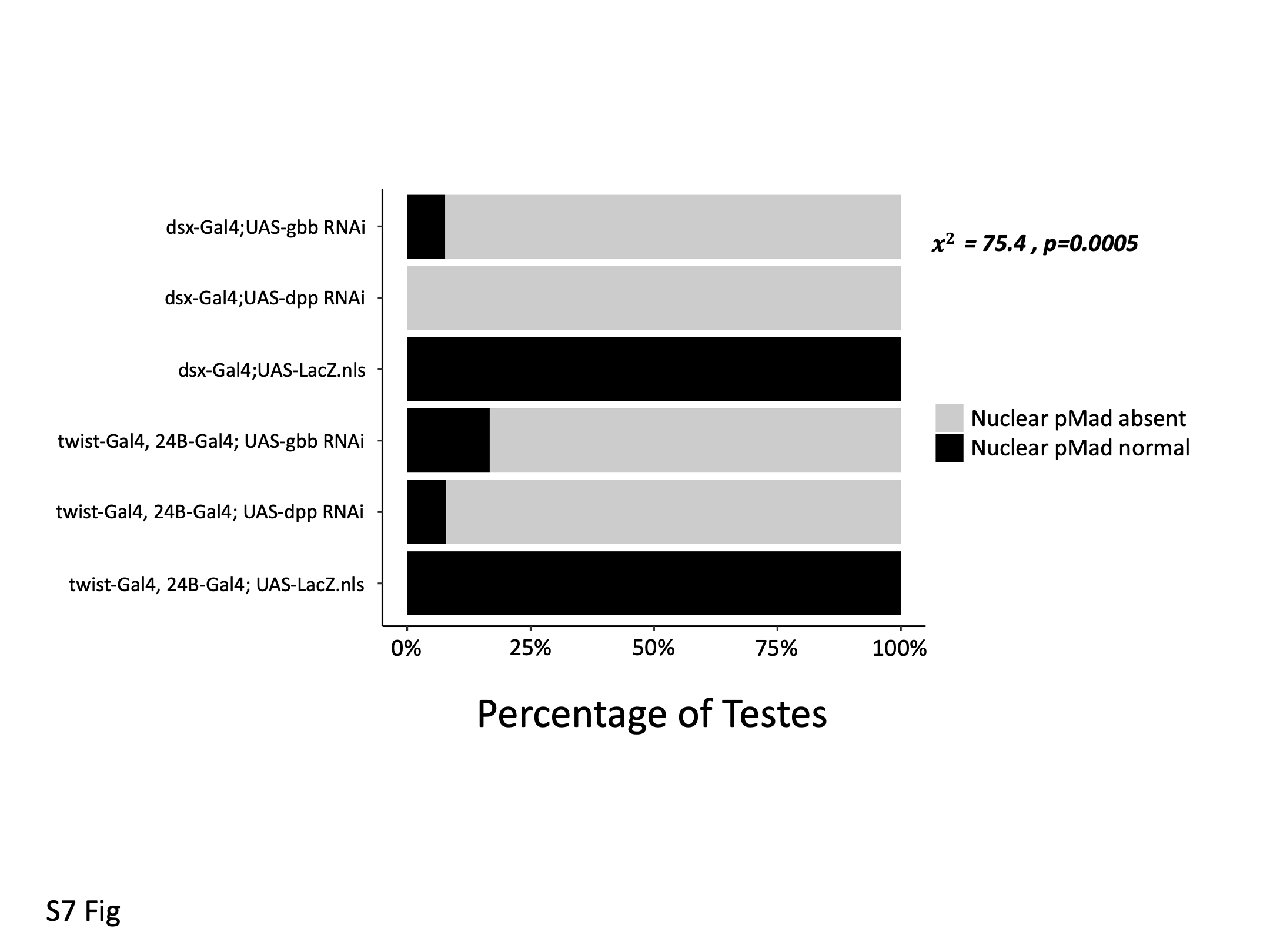

### Supplemental Fig8

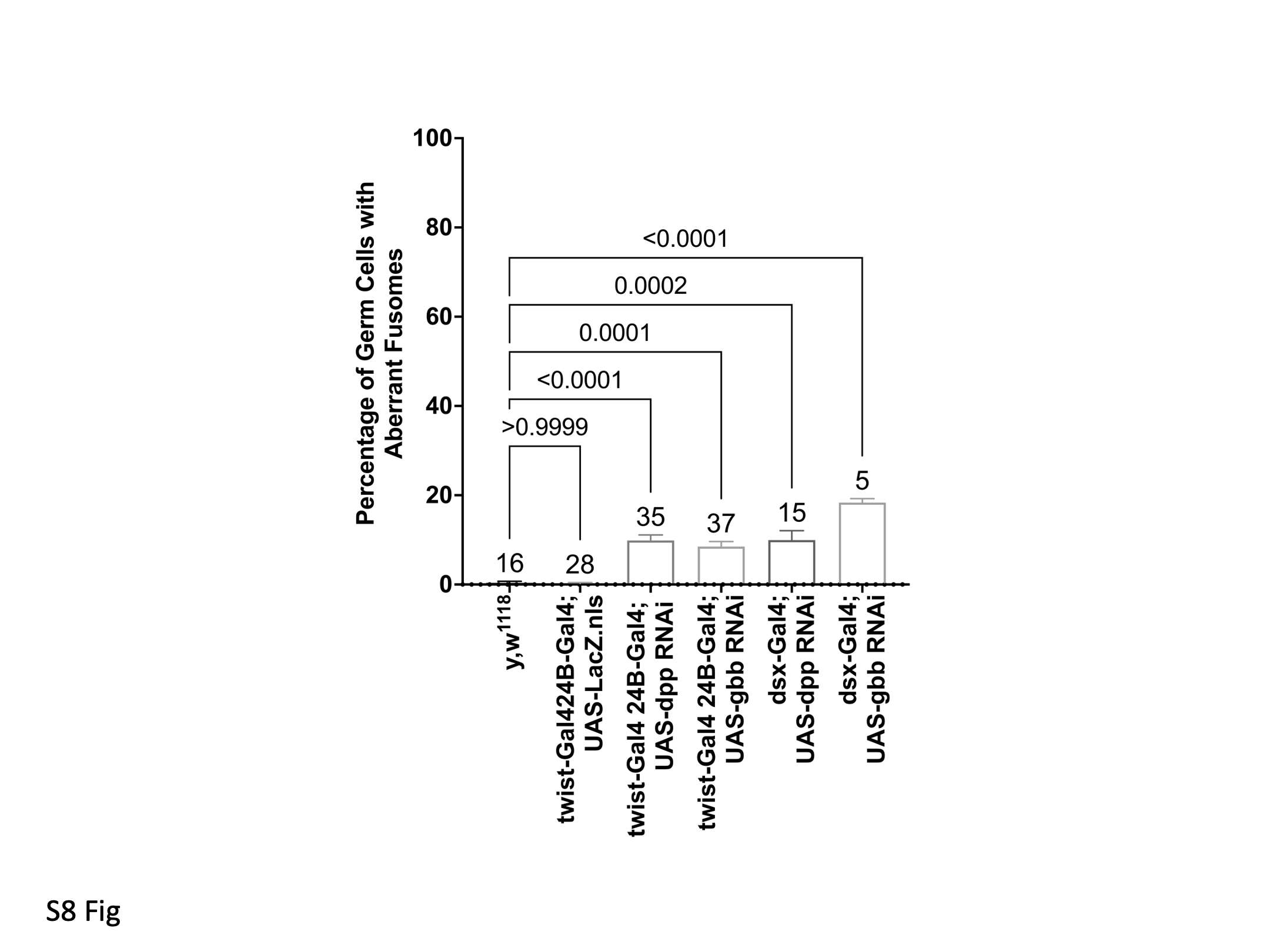
